# *In vitro* Efficacy of Novel Glucan Synthase Inhibitor, Ibrexafungerp (SCY-078), Against Multidrug- and Pan-resistant *Candida auris* Isolates from the Outbreak in New York

**DOI:** 10.1101/811182

**Authors:** Yan Chun Zhu, Stephen A. Barat, Katyna Borroto-Esoda, David Angulo, Sudha Chaturvedi, Vishnu Chaturvedi

**Affiliations:** Mycology Laboratory, Wadsworth Center, New York State Department of Health, Albany, NY, USA; SCYNEXIS, Inc., Jersey City, NJ, USA

## Abstract

We report low MIC_50_ of Ibrexafungerp (SCY-078) for 102 *Candida auris* clinical and surveillance isolates from outbreak in New York. The group included *C. auris* with a variable resistance to antifungal drugs. Five pan-resistant *C. auris* isolates were susceptible to Ibrexafungerp with low MIC_50_ range of 0.12-1 µg/ml.

## Introduction

Since its discovery in 2009, *Candida auris* has become a major concern as an emerging drug resistant, healthcare-related infection around the globe (1, 2). Beginning in 2013, the New York metropolitan area has suffered from a large, sustained outbreak of *C. auris* in hospitals and healthcare facilities (3). The latest NY data from 540 *C. auris* clinical isolates, *C. auris* isolates recovered from 11,035 patient surveillance specimens, and 3,672 environmental surveillance samples suggests the predominance of the South Asia Clade I with variable multidrug-resistance (4). There is a pressing public health need to test additional antifungal drugs for their efficacy against drug resistant *C. auris*.

Ibrexafungerp (formerly SCY-078), is an orally bioavailable, semi-synthetic modified compound of enfumafungin, a triterpene glycoside natural product (5, 6). Ibrexafungerp laboratory testing showed broad activity against *Candida* species including fluconazole-resistant *C. albicans*, and *C. auris* (7-9). We report *in vitro* activity of Ibrexafungerp against 102 *C. auris* isolates from NY comprising clinical and surveillance cases. The *C. auris* selection included variable resistance (resistance to one drug in one or two classes of antifungal drug) multidrug-resistant (resistance to two or more drugs between two classes of antifungal drugs) or pan-resistant (resistance to two or more azoles, all echinocandins, and amphotericin B).

## Materials and Methods

A recent publication from our laboratory includes details about the processing, identification, and characterization of *C. auris* from the NY outbreak (4). Broth micro-dilution antifungal susceptibility testing was performed per Clinical and Laboratory Standards Institute reference method M27-A3 as described by Berkow et al. (7). Amphotericin B MIC_100_ was tested using E-test strips (bioMerieux USA, St. Louis, MO). We used CDC guidelines to assess antifungal resistance patterns in *C. auris* as breakpoints are not available for this pathogen (https://www.cdc.gov/fungal/candida-auris/c-auris-antifungal.html). We also used EUCAST recommendations as a surrogate breakpoint to determine resistance pattern to various antifungals (http://www.eucast.org/fileadmin/src/media/PDFs/EUCAST_files/AFST/Clinical_breakpoints/Antifungal_breakpoints_v_9.0_180212.pdf).

## Results and Discussion

Ibrexafungerp MIC_50_ readings for 102 *C. auris* isolates are summarized in Tables 1-3. For 97 *C. auris* isolates with a variable resistance to antifungal drugs, Ibrexafungerp MIC_50_ range was 0.06-0.5 µg/ml, median and mode were 0.5 µg/ml, respectively (Table 1-2). All five pan-resistant *C. auris* isolates were susceptible to Ibrexafungerp with low MIC_50_ range of 0.12-1 (Table 3). Notably, Ibrexafungerp MIC_50_ range against our collection is well within the serum achievable concentrations reported from preclinical pharmacokinetics and pharmacodynamic studies, and murine models of disseminated candidiasis (10). Our laboratory findings expand earlier reports on *in vitro* efficacy of Ibrexafungerp against *C. auris* resistant to amphotericin B, flucytosine, itraconazole and isavuconazole (7, 9). The near universal fluconazole resistance, and unusually high resistance to azoles, echinocandins, polyene, and nucleoside inhibitors is a noticeable feature of *C. auris* isolates in the NY outbreak (3, 4). The exact mechanisms behind the observed resistance pattern remain unknown; with the exception of few isolates, all NY *C. auris* isolates belonged to South Asia clade, which is reported to have high drug-resistance (2). Our findings support enhanced evaluations of Ibrexafungerp including expanded clinical studies to better understand its therapeutic potential for *C. auris*.

**Table 1.**
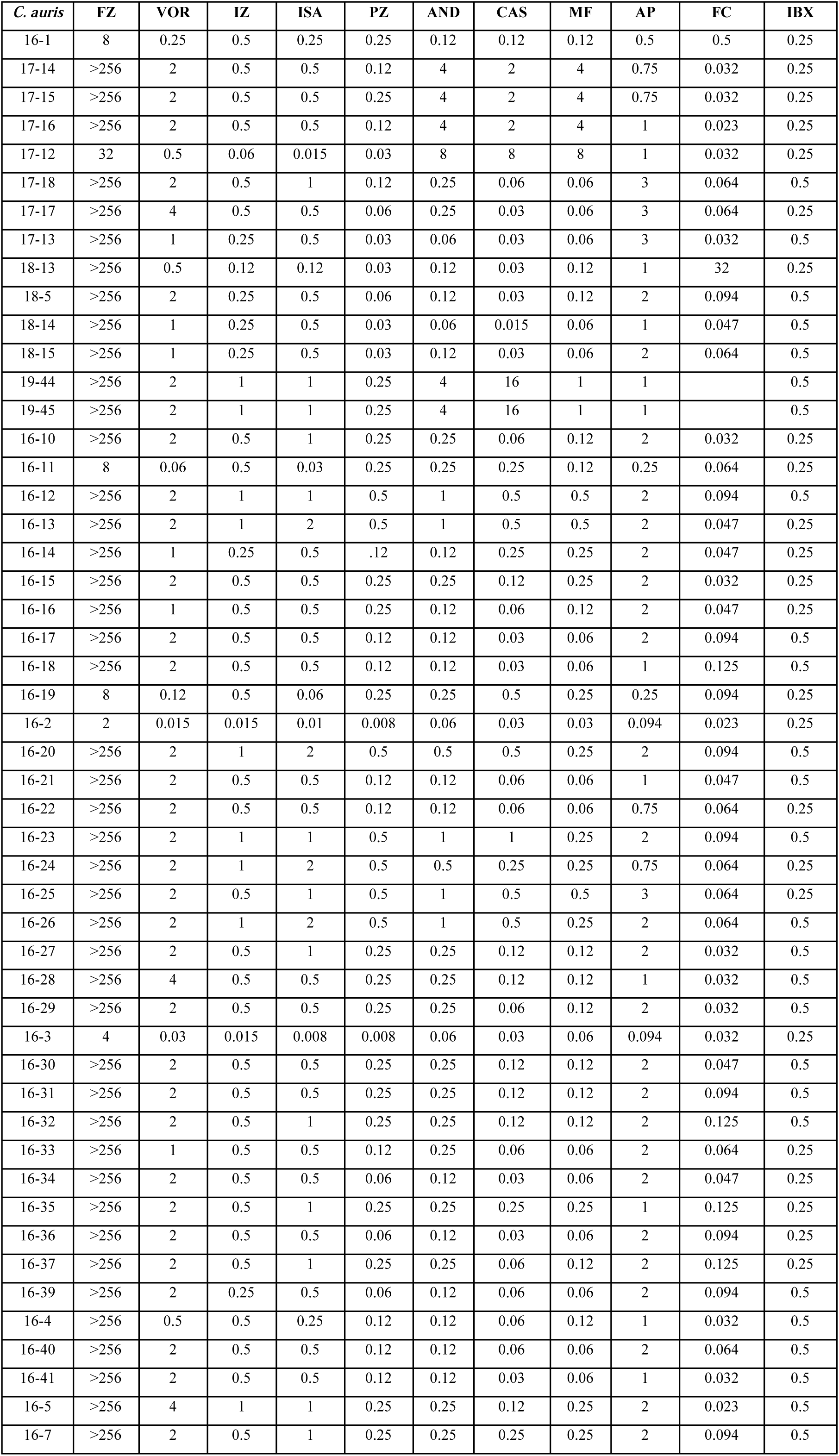
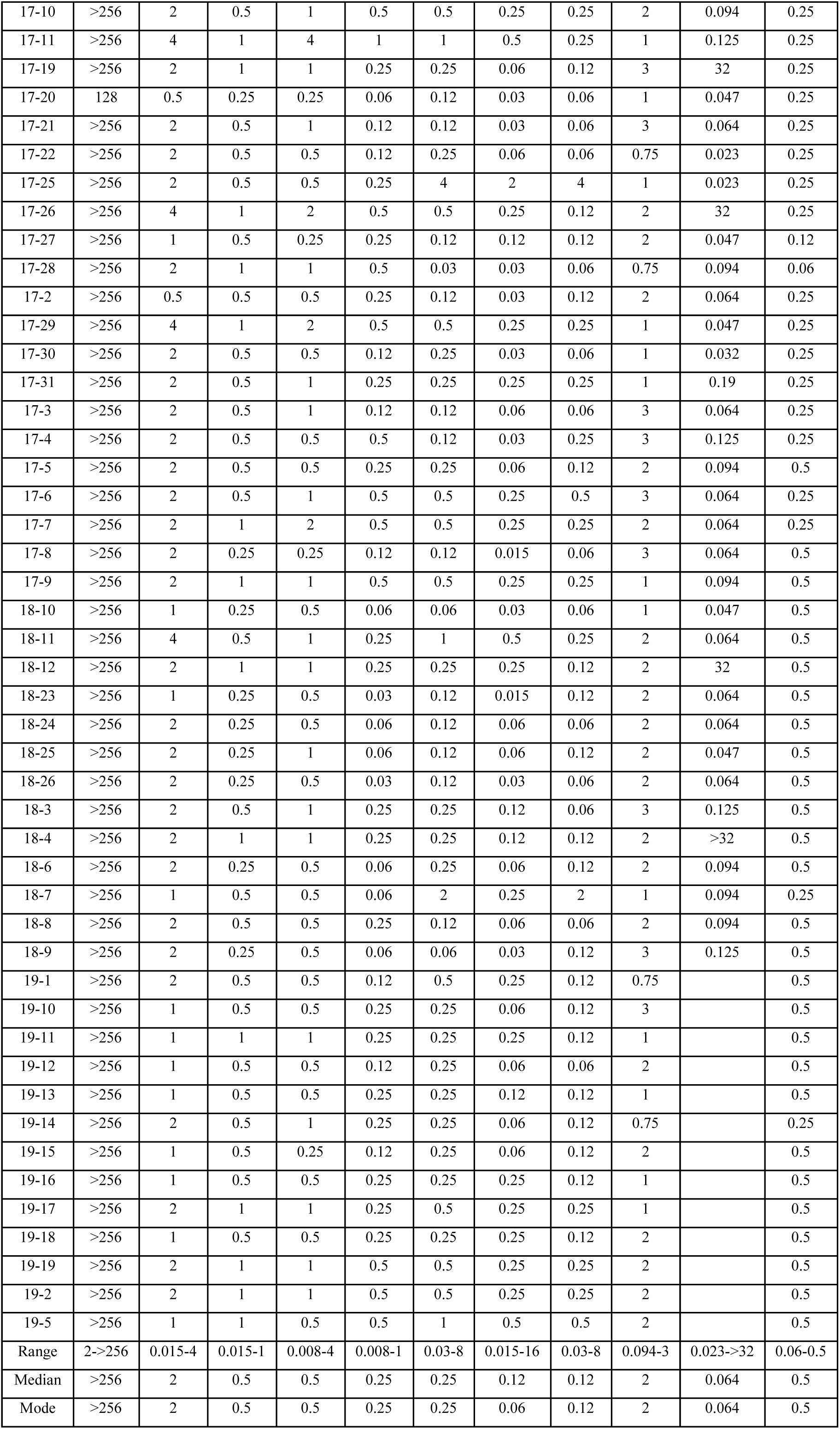
Detailed Ibrexafungerp MIC_50_ (µg/ml) for 98 clinical and surveillance isolates of *Candida auris*

**Table 2.** Summary Ibrexafungerp MIC_50_ (µg/ml) for 98 clinical and surveillance isolates of *Candida auris*

| FZ |  |  | VOR |  |  | IZ |  |  | ISA |  |  | PZ |  |  | AND |  |  | CAS |  |  | MF |  |  | AP |  |  | FC |  |  | IBX |  |  |
| --- | --- | --- | --- | --- | --- | --- | --- | --- | --- | --- | --- | --- | --- | --- | --- | --- | --- | --- | --- | --- | --- | --- | --- | --- | --- | --- | --- | --- | --- | --- | --- | --- |
| Range | Median | Mode | Range | Median | Mode | Range | Median | Mode | Range | Median | Mode | Range | Median | Mode | Range | Median | Mode | Range | Median | Mode | Range | Median | Mode | Range | Median | Mode | Range | Median | Mode | Range | Median | Mode |
| 2 -<br>>256 | >256 | >256 | 0.015<br>- 4 | 2 | 2 | 0.015<br>- 1 | 0.5 | 0.5 | 0.008<br>- 4 | 0.5 | 0.5 | 0.008<br>- 1 | 0.25 | 0.25 | 0.03 -<br>8 | 0.25 | 0.25 | 0.015<br>- 16 | 0.12 | 0.06 | 0.03 -<br>8 | 0.12 | 0.12 | 0.094<br>- 3 | 2 | 2 | 0.023<br>- >32 | 0.064 | 0.064 | 0.06 –<br>0.5 | 0.5 | 0.5 |

**Table 3.** Ibrexafungerp MIC_50_ data for pan-resistant *Candida auris*

| *Candida auris* | MIC<sub>50</sub> |  |  |  |  |  |  |  |  |  |  |
| --- | --- | --- | --- | --- | --- | --- | --- | --- | --- | --- | --- |
|  | FZ | VOR | IZ | ISA | PZ | AND | CAS | MF | AP | FC | IBX |
| **17-24<sup>#</sup>** | **>256** | **2** | **0.5** | **0.5** | **0.25** | **4** | **>16** | **4** | **2** | **0.064** | **0.5** |
| **18-2<sup>#</sup>** | **>256** | **2** | **1** | **1** | **0.25** | **8** | **2** | **4** | **2** | **0.064** | **0.12** |
| **19-4<sup>#</sup>** | **>256** | **2** | **0.5** | **0.5** | **0.25** | **4** | **2** | **4** | **2** | **NA** | **0.5** |
| **19-42<sup>#</sup>** | **>256** | **2** | **1** | **1** | **0.25** | **4** | **16** | **4** | **2** | **NA** | **1** |
| **19-43<sup>#</sup>** | **>256** | **2** | **1** | **1** | **0.25** | **4** | **16** | **4** | **2** | **NA** | **1** |

<sup>#</sup>**Pan-resistant**
AP (E-test)
NA not available

## ACKNOWLEDGEMENTS

We thank Brittany O’Brien and Jiali Liang for their contributions to antifungal susceptibility testing at the Mycology Laboratory, Wadsworth Center; Thomas Chen, SCNYNEXIS, Inc., is thanked for the editorial comments. Stephen A. Barat, Katyna Borroto-Esoda, and David Angulo are employed by SCYNEXIS, Inc., the manufacturer of Ibrexafungerp.

## References

1. Satoh K, Makimura K, Hasumi Y, Nishiyama Y, Uchida K, Yamaguchi H. 2009. *Candida auris sp. nov*., a novel ascomycetous yeast isolated from the external ear canal of an inpatient in a Japanese hospital. Microbiology and immunology 53:41–44.

2. Lockhart SR, Etienne KA, Vallabhaneni S, Farooqi J, Chowdhary A, Govender NP, Colombo AL, Calvo B, Cuomo CA, Desjardins CA. 2017. Simultaneous Emergence of Multidrug-Resistant *Candida auris* on 3 Continents Confirmed by Whole-Genome Sequencing and Epidemiological Analyses. Clinical Infectious Diseases 64:134–140.

3. Adams E, Quinn M, Tsay S, Poirot E, Chaturvedi S, Southwick K, Greenko J, Fernandez R, Kallen A, Vallabhaneni S, Haley V, Hutton B, Blog D, Lutterloh E, Zucker H, Workgroup. CaI. 2018. *Candida auris* in Healthcare Facilities, New York, USA, 2013-2017. Emerging Infectious Diseases 24:1816–1824.

4. Zhu Y, O’Brien B, Leach L, Clark A, Bates M, Adams E, Ostrowsky B, Quinn M, Dufort E, Southwick K, Erazo R, Haley VB, Bucher C, Chaturvedi V, Limberger RJ, Blog D, Lutterloh E, Chaturvedi S. 2019. Laboratory Analysis of an Outbreak of *Candida auris* in New York from 2016 to 2018-Impact and Lessons Learned. bioRxiv doi:10.1101/760090:760090.

5. Schwartz RE, Smith SK, Onishi JC, Meinz M, Kurtz M, Giacobbe RA, Wilson KE, Liesch J, Zink D, Horn W, Morris S, Cabello A, Vicente F. 2000. Isolation and Structural Determination of Enfumafungin, a Triterpene Glycoside Antifungal Agent That Is a Specific Inhibitor of Glucan Synthesis. Journal of the American Chemical Society 122:4882–4886.

6. Apgar JM, Wilkening RR, Greenlee ML, Balkovec JM, Flattery AM, Abruzzo GK, Galgoci AM, Giacobbe RA, Gill CJ, Hsu MJ, Liberator P, Misura AS, Motyl M, Nielsen Kahn J, Powles M, Racine F, Dragovic J, Habulihaz B, Fan W, Kirwan R, Lee S, Liu H, Mamai A, Nelson K, Peel M. 2015. Novel orally active inhibitors of β-1,3-glucan synthesis derived from enfumafungin. Bioorganic & Medicinal Chemistry Letters 25:5813–5818.

7. Berkow EL, Angulo D, Lockhart SR. 2017. In Vitro Activity of a Novel Glucan Synthase Inhibitor, SCY-078, against Clinical Isolates of *Candida auris*. Antimicrobial Agents and Chemotherapy 61:e00435–17.

8. Borroto-Esoda K, Barat S, Angulo D, Sobel R. 2019. In Vitro Activity of Ibrexafungerp (SCY-078) Against *Candida* spp. (Including Fluconazole-Resistant Isolates) [14OP]. Obstetrics & Gynecology 133:S1.

9. Larkin E, Hager C, Chandra J, Mukherjee PK, Retuerto M, Salem I, Long L, Isham N, Kovanda L, Borroto-Esoda K, Wring S, Angulo D, Ghannoum M. 2017. The Emerging Pathogen *Candida auris*: Growth Phenotype, Virulence Factors, Activity of Antifungals, and Effect of SCY-078, a Novel Glucan Synthesis Inhibitor, on Growth Morphology and Biofilm Formation. Antimicrobial Agents and Chemotherapy 61:e02396–16.

10. Wring SA, Randolph R, Park S, Abruzzo G, Chen Q, Flattery A, Garrett G, Peel M, Outcalt R, Powell K, Trucksis M, Angulo D, Borroto-Esoda K. 2017. Preclinical Pharmacokinetics and Pharmacodynamic Target of SCY-078, a First-in-Class Orally Active Antifungal Glucan Synthesis Inhibitor, in Murine Models of Disseminated Candidiasis. Antimicrobial Agents and Chemotherapy 61:e02068–16.

